# *Chlamydia trachomatis* infection upregulates MicroRNA21 to deplete tumor suppressor PTEN

**DOI:** 10.64898/2026.05.11.723860

**Authors:** Karthika Rajeeve, Suvagata Roy Chowdhury, Marco Albrecht, Nadine Vollmuth, Jörg Wischhusen, Thomas Rudel

## Abstract

Infection with obligate intracellular *Chlamydia trachomatis* (Ct) has been associated with cervical and ovarian carcinoma in humans. The possible cause of the tumor-promoting effect of the infection is thought to be a manipulation of host signaling pathways, which are essential for the growth of the bacteria, and which at the same time can support tumor growth. The PI3K and MAPK signaling cascades are outstanding candidates for this concept, as they are persistently activated during infection and required for the growth of *Chlamydia* and many tumors. The mechanism by which *Chlamydia* activates these pre-transforming survival signals in the cells is not well understood. Here we show that Ct infection up-regulates the oncomiR, miR21 by activating the host transcription factor AP-1. This depletes the target gene of miR-21, the tumor suppressor PTEN that ensures the persistent activation of the PI3K pathway. Blocking miR21 adversely affects the growth and development of the pathogen. We show here that miR21 KO mice are less susceptible to infection with Ct compared to control mice. Our data thus provides direct *in vivo* evidence of the induction and dependency of this obligate human pathogen on tumor-promoting miR-21-induced signaling.

**Importance:** *Chlamydia trachomatis* is an obligate intracellular pathogen that relies on host metabolism for survival, yet the mechanisms by which it sustains nutrient access remain incompletely understood. Here, we identify a conserved host regulatory axis in which infection induces miR-21 to deplete the tumor suppressor PTEN, thereby enabling persistent activation of PI3K signaling. This pathway promotes a tumor-like metabolic state that supports bacterial growth while protecting infected cells from apoptosis. We further demonstrate that the AP-1-miR-21-PTEN circuit is required for efficient chlamydial replication *in vitro* and *in vivo*, and that disruption of miR-21 significantly impairs bacterial propagation in epithelial tissues. These findings reveal how *C. trachomatis* hijacks a central oncogenic signaling network to remodel host cell physiology and highlight a mechanistic link between infection and cancer-associated pathways. Targeting this host-driven signaling axis may provide new strategies for controlling infection independent of traditional antibiotics.

## Introduction

Sexually transmitted diseases are a major economic burden and have a long-term impact on health. Of the 357 million new cases of sexually transmitted infections each year, 131 million cases are due to *Chlamydia trachomatis* (Ct) making it the most common cause of bacterial sexually transmitted infection (1). Ct infections are frequently asymptomatic; transmission and re-infection occur often even after antibiotic therapy. Chronic and persistent infection leads to inflammatory pathology like the development of pelvic inflammatory diseases, scaring of the infected tissue, and eventually sterility. In addition, *Chlamydia* infection has been closely associated with the development of ovarian and cervical cancer (2, 3). The molecular evidence of this etiology is not yet known. We and other groups have previously reported that *Chlamydia* infection leads to depletion of the tumor suppressor p53 (4, 5) and DNA double-strand breaks (6). The infection also stabilizes the proto-oncogene c-Myc in the infected tissue, another possible link of the infection-cancer interface (7, 8).

*Chlamydia* is an obligate intracellular human pathogen with a biphasic life cycle. The infectious form of the bacteria is the elementary body (EB), which upon entering the cells forms a membrane-bound inclusion where the EB transforms into the replicative reticulate body (RB). After several cycles of replication, the RBs switch back to EBs, which are released into the extracellular milieu from where they start a new infection cycle. Since *Chlamydia* leads a parasitic lifestyle, it modulates and rewires host signaling pathways to adapt to its niche. The bacterium with a reduced genome is highly dependent on the uptake of host metabolites (9).

To meet their metabolic needs, the bacteria have been shown to activate various host signaling pathways (10, 11). TP53 and c-Myc, which are regulated during Ct infection, are well-known tumor-associated proteins that play a crucial role in maintaining the metabolic pathways in the host cells (12, 13). The most important upstream signaling pathway regulating all these metabolic processes is the PI3K-AKT signaling network, which is activated via receptor tyrosine kinases (RTK), G protein-coupled receptors, integrins, and cytokine receptors (14). Chlamydial infection is known to activate the PI3K and MAPK signaling pathway immediately after infection and throughout its developmental cycle and is critical for the growth and establishment of the pathogen (15, 16). Little is known about how the pathogen maintains the signaling pathway throughout its life cycle. Chlamydial infection has been shown to activate PI3K via the EphA2 receptor (15). Furthermore, the virulence factor of *Chlamydia,* TepP is secreted early in infection to spatially restrict the activation of the PI3K pathway (16).

MicroRNAs (miRNAs) are small noncoding RNAs (ca. 20-22 nt) expressed by eukaryotic cells which play an important role in host-bacteria interaction (17). It is estimated that these single-stranded RNAs regulate up to 60% of the genes in humans. The effective miRNA is processed from larger transcripts and forms a miRNA duplex. One strand of the duplex is then incorporated into the miRNA-induced silencing complex (miRISC), which represses target mRNA function by a combination of translational repression and mRNA degradation (18). The deregulation of miRNAs is a crucial response of the host to invading microbes. At the same time, the pathogens manipulate these macromolecules to reprogram the host cells for their purposes. In the context of chlamydial infection, several reports suggest an important role of miRNAs during infection of humans and mice (19–23). These potential biomarkers are deregulated in endo-cervical samples of women infected with *C. trachomatis* (24). In mice, Ct infection has been shown to lead to depletion of miR214, resulting in upregulation of ICAM 1, which contributes to hydrosalpinx and upper genital tract pathology (25). Interestingly, several miRNAs were connected to conjunctival inflammation and ocular scarring (26). Immune modulatory functions have been attributed to miR378b and miR155, controlling genital tract pathology and anti-chlamydial immunity, respectively (21, 27). Previous studies in our laboratory have shown that *Chlamydia* infection induces a plethora of changes in miRNA regulation in infected host cell (19), including the up-regulation of miR30c-5p, resulting in depletion of TP53 and DRP1, which prevents mitochondrial fission and thus preserves their integrity (19).

Here, we explore another signaling pathway that *Chlamydia* redirects in the host and reveal its association with malignancy. We show that Ct infection upregulates miR21, a member of the oncomiR family that acts as a tumor suppressor and oncogene (28, 29). This leads to the depletion of the tumor suppressor PTEN, which acts as a potent inhibitor of PI3K. In addition, miR21 KO mice do not support Ct growth in the female genital tract suggesting a dependency of *Chlamydia* on miR21 signaling pathways *in vivo*.

## Results

### *C. trachomatis* infection leads to depletion of the tumor suppressor PTEN

*C. trachomatis* is known to activate the MAPK and PI3K signaling pathways in epithelial cells (15), for example in infected HeLa229 cells (Fig. 1A). To understand if this deregulated survival signal during chlamydial infection also applies to primary cells, we extracted primary cells from human fimbriae (Fimb cells). The chlamydial infection of Fimb cells showed larger inclusions than that of HeLa cells and these infected cells produced infectious progeny (data not shown). Since infection induces persistent activation of these survival pathways, we investigated if the tumor suppressor PTEN which is a negative regulator of the PI3K pathway is affected during Ct infection. Western blot analysis of Fimb cells infected with the serovar Ct L2 and D at different time points showed an infection-induced depletion of PTEN levels (Fig. 1B,C). To confirm that this is not a cell type-and species-dependent effect, we used mouse fimbriae cells and primary MEFs for the study. Infection with *C. trachomatis* and *C. muridarum* led to a decrease in PTEN levels in different cell types (Fig. 1D, S1A,B). To understand the mechanism by which Ct affects PTEN protein levels we isolated total RNA from the infected cells and analyzed PTEN mRNA levels by RT-PCR. Interestingly, the mRNA level of PTEN was significantly reduced after Ct infection (Fig. 1E), suggesting a possible role of miRNA in the regulation of PTEN.

**Figure 1.**
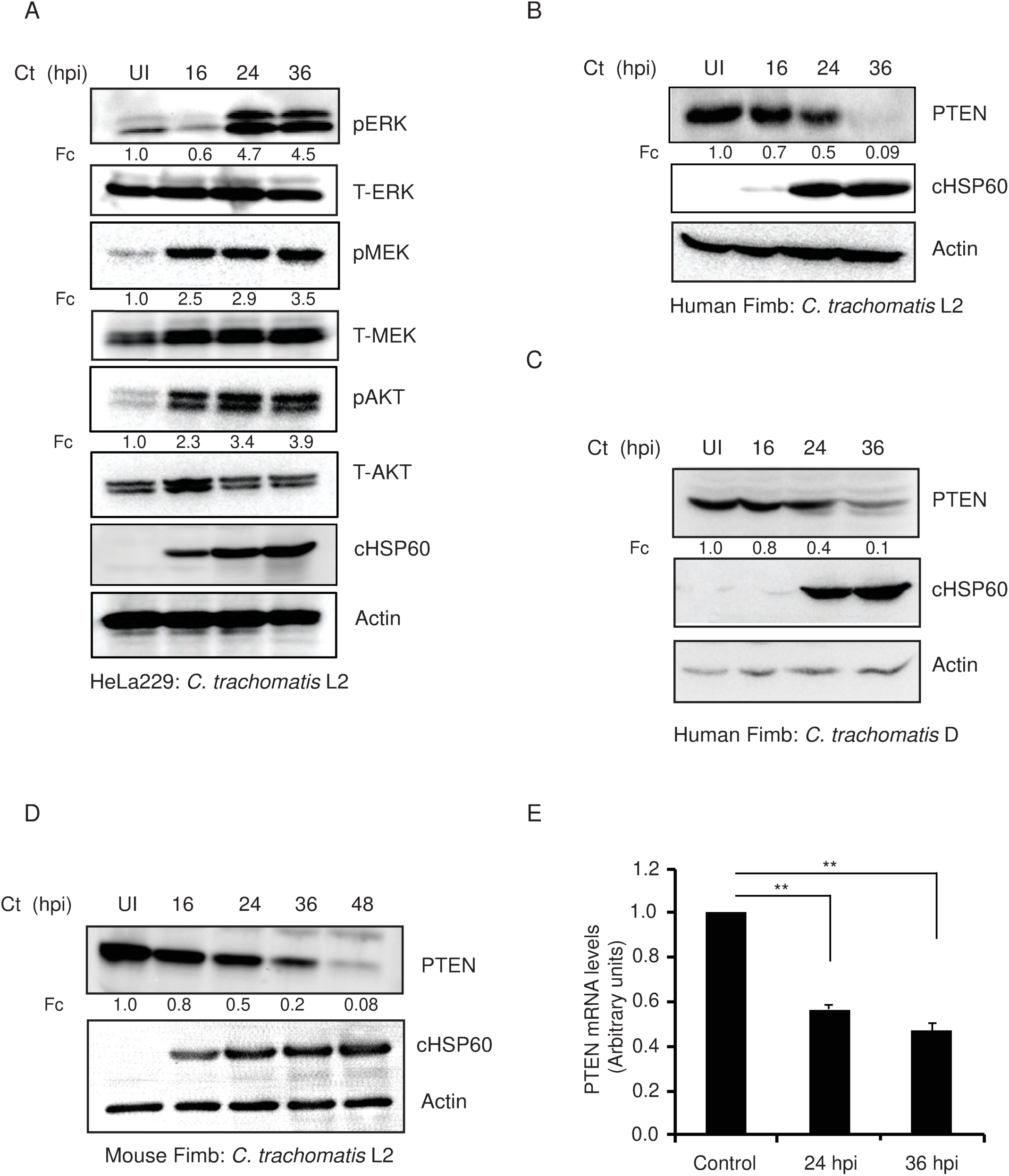
*Chlamydia trachomatis* infection leads to the depletion of the tumor suppressor PTEN. **A**. HeLa229 cells were infected with Ct L2 serovar at a MOI of 1 for different time points; the cells were lysed and analyzed via western blotting to detect PI3K and MAPK signaling pathways. n=3 **B**. Cells isolated from human fimbriae (human Fimb) were infected with Ct L2 serovar at a MOI of 1 for different time points; the cells were lysed and analyzed via western blotting to detect levels of PTEN. n=3. **C.** Human Fimbs were infected with Ct D serovar at a MOI of 1 for various time points. The lysate was probed against PTEN antibody. n=3 **D**. Mouse Fimb cells were infected with Ct L2 serovar at a MOI of 1 for different time points to detect levels of PTEN after western blotting. For the above experiments n=3 Beta actin serves as the loading control and cHSP60, Chlamydial HSP60, detects the rate of infection, UI= uninfected. Fc shows the fold change in the levels of protein compared to UI, normalized to levels of beta actin. **E.** Quantitative RT-PCR to detect PTEN mRNA levels in HeLa229 cells infected with Ct L2. There independent experiments were performed with the mean values (±SEM) compared to levels of uninfected HeLa229 cells. n=3. Statistical analysis was performed using Student t test (**p ≤ 0.01).

### *Chlamydia trachomatis* infection activates miR21 to deplete its target PTEN

It is well known that miRNAs play an important role in the downregulation of PTEN in tumor cells (30, 31). We previously performed a screen for differentially expressed miRNAs in *C. trachomatis-*infected primary human umbilical vein endothelial cells (HUVECs) (19). Of the number of miRNAs deregulated during Ct infection one of the prominent miRNAs known to regulate PTEN was miR21. miRSeq showed a 2.4-times upregulation of miR21 upon Ct infection. This was confirmed in HeLa cells after Ct infection by RT PCR quantifying the mature miR21 levels (Fig. 2A, D). We also analyzed miR21 levels after Ct infection after different time points of infection using Northern blotting in HeLa229, HUVECS, and Fimb cells (Fig. 2B,C). Interestingly heat killed bacteria failed to activate miR21 expression (Fig. 2A), suggesting that Chlamydia actively manipulates miR21 levels in these cells.

**Figure 2.**
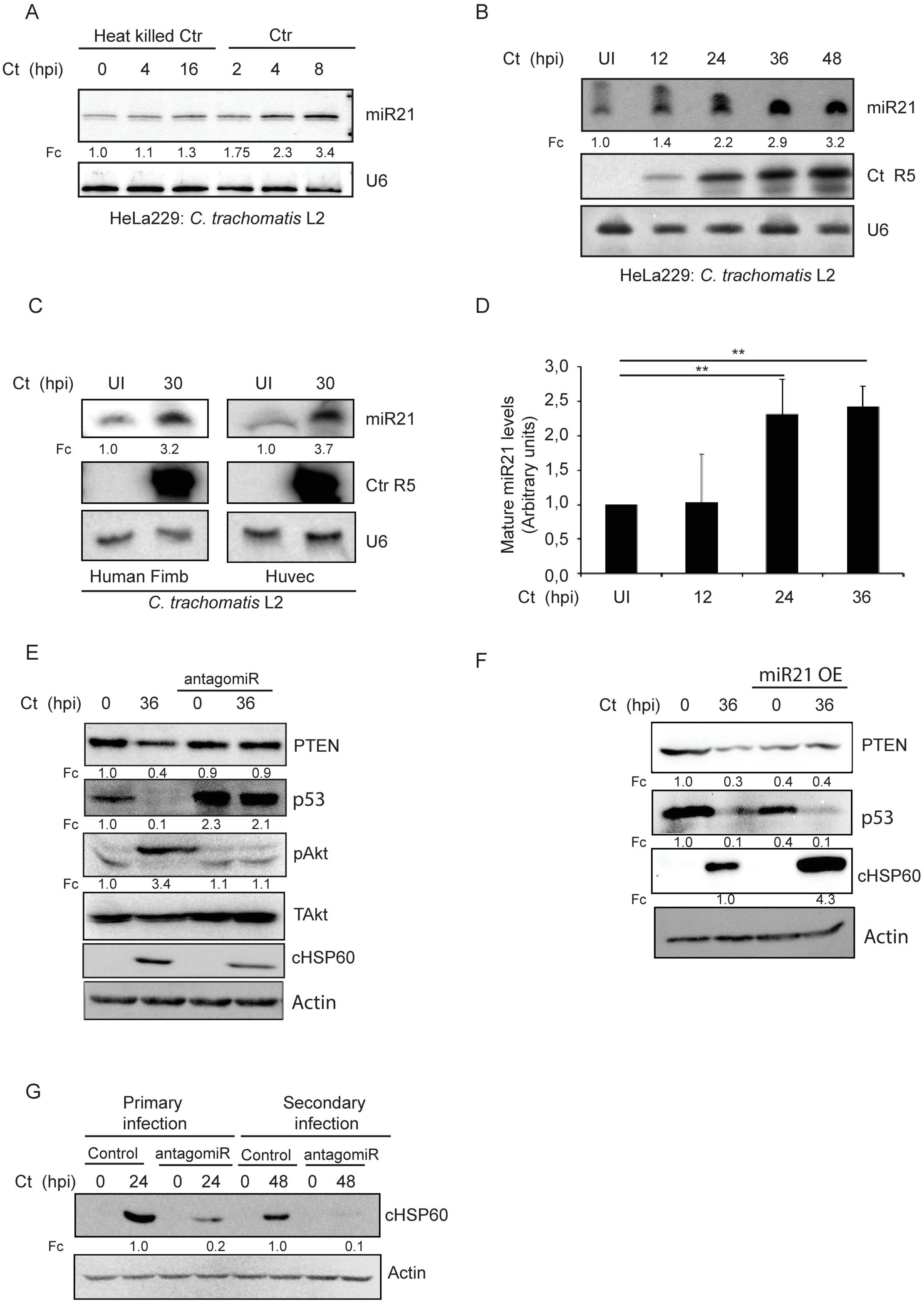
*Chlamydia trachomatis* infection leads to up regulation of oncomiR, miR21 to deplete PTEN level. **A.** Hela229 cells were infected with heat killed or infectious Ct for different (early) time points and miR21 levels were detected by northern blotting. U6 was detected as loading control. **B.** Hela229 cells were left uninfected (UI) or infected with infectious Ct for different time points. The total RNA was used to perform northern blotting to detect levels of miR21. U6 serves as loading control and Ct R5 is the infection control. n=3. **C.** Human Fimb and Huvec cells were infected with infectious Ct for 30 h and the total RNA was used for Northern blot analysis to detect the levels of miR21; UI-uninfected, Ct R5-infection control and U6 is the loading control. **D.** HeLa229 cells were either left uninfected or infected with Ct for different time points. The total RNA was isolated and quantitative RT-PCR was used to quantify the levels of miR21. Three independent experiments were performed with the mean values (±SEM) compared to levels of uninfected HeLa229 cells. Statistical analysis was performed using Student *t* test (**p ≤ 0.01). **E.** HeLa229 cells were treated with scrambled RNA or antagomiR against miR21 or **F** over expressed with scramble RNA or miR21 RNA. The cells were left uninfected or infected with Ct for 36h and lysed for western blot analysis. The data provided is the representative of three independent experiments. n=3. **G.** The cells (uninfected and infected) from **E** (primary infection) were lysed to re-infect freshly plated HeLa229 cells to analyze the progeny, secondary infection. For the above experiments **F**. and **G**. Beta actin serves as the loading control and cHSP60, Chlamydial HSP60, detects the rate of infection. Fc shows the fold change in the levels of protein compared to UI, normalized to levels of beta actin. The data provided is the representative of three independent experiments. n=3.

### Downregulation of miR21 restricts the growth of *C. trachomatis*

To investigate if miR21-dependent PTEN regulation and downstream signaling via PI3K is critical for chlamydial growth, we treated *Chlamydia* infected and uninfected HeLa229 cells with an antagomir, inhibitor against miR21 or scrambled oligonucleotide to analyze the regulation of PTEN and other downstream signals via Western blotting. Blocking miR21 using antagomir not only stabilized the level of PTEN but also TP53, resulting in the inactivation of pAkt (Fig. 2E). This also restricted the growth and development of the bacteria as evident from the progeny analysis (Fig. 2G). Interestingly, overexpression of miR21 had the opposite and positive effect on bacterial growth, reducing levels of PTEN and P53 with a corresponding 4.3-fold increase in *Chlamydia* growth (Fig. 2F). These data showed that miR21 is upstream of the PI3K signaling pathway and plays a pivotal role in the development of the pathogen.

To investigate the infection-induced miR21-upregulation in intact HeLa229 cells, we made constructs with miR21 targeting sites fused to GFP. To detect dynamic changes in GFP levels, a PEST sequence was fused to GFP to reduce the stability and half-life of the protein (32). This enables a rapid loss of signal with miR21 binding (Fig. S2). In addition, a dsRED expression cassette was included in this construct to detect transfected cells independent of GFP levels (Fig. 3A). Only cells expressing the miR21 PEST-GFP construct, but not those without miR21 targeting sequence (PEST-GFP) lost the GFP signal upon infection with *Chlamydia*, consistent with the upregulation of miR21 in infected cells and specific downregulation of the miR21 PEST-GFP (Fig. 3A,B,C,D). We then cloned the 3’ - untranslated region of *PTEN* (PUG) with a natural miR21 binding into the PEST GFP construct to test if infection-induced upregulated miR21 can directly target the PTEN sequence. In addition, we introduced a point mutation into the binding site of the miR21 targeting sequence (M-PUG) and fused it to PEST GFP. Expression of PUG but not M-PUG resulted in the loss of the GFP signal (Fig. 3E,F) during *Chlamydia* infection, consistent with the direct targeting of PTEN by the upregulation of miR21.

**Figure 3.**
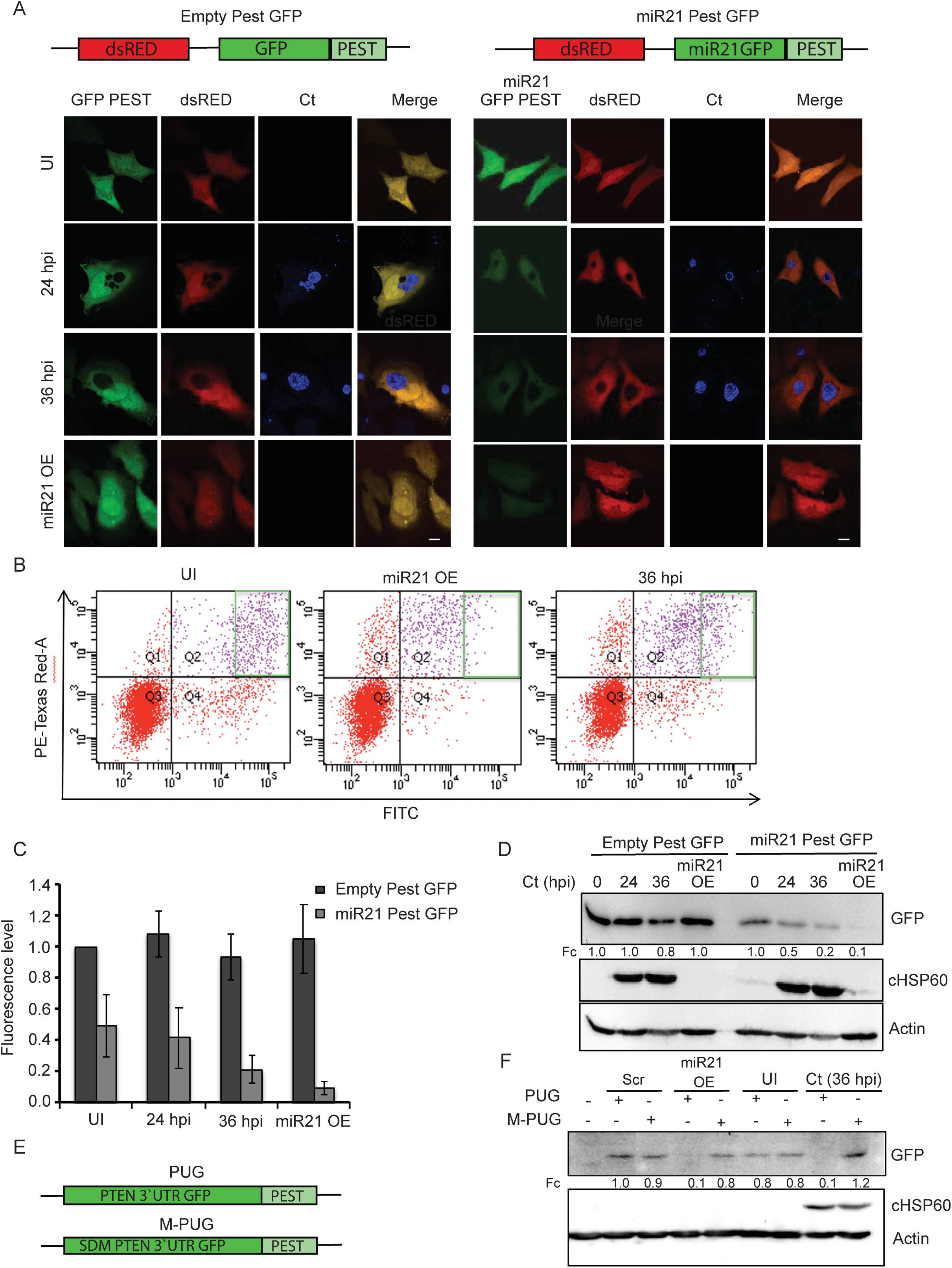
miR21 is up-regulated in *Chlamydia trachomatis* infected cells. **A.** PEST-GFP construct with empty or miR21 binding site were constructed. dsRed is used as control to detect transfection efficiency. These cells were with left uninfected (UI) or infected with Ct (for 24h, 36h), or over expressed with miR21. The cells were fixed and analyzed using immunostaining. Green indicates GFP signal, red-dsRED, and blue-Chlamydial inclusion. The images represent three independent experiments. **B.** The cells from **A** were subjected to FACS analysis. The distribution of red (PE-Texas Red) and green (FITC) cells are depicted in the green gate given. **C**. The mean fluorescence value from FACS data from **B** is quantified and presented as graph. **D**. The cells from **B** were taken for western blot analysis to detect the levels of GFP. Beta actin and cHSP60 serve as a leading control and infection level indicator respectively. **E**. The 3 prime UTR of PTEN, the miR21 binding target (PUG) and the site directed mutated 3 prime UTR of PTEN (M-PUG) was cloned into the PEST GFP **F.** These cells were either transfected with scrambled peptide or miR21 OE and left uninfected (UI) or infected with Ct. The cells were lysed to perform western blot analysis to detect levels of GFP. cHSP60 serve to detect Ct infection and beta actin serve as loading control. All the experiments were repeated thrice (n=3).

### *Chlamydia* infection activates the transcription factor AP-1 to up-regulate miR21

To investigate how chlamydial infection induces upregulation of miR21, we further explored the mechanism. It is known that activating protein-1 (AP-1) drives miR21 transcription. Ap-1 consists of DNA-binding proteins of the Jun, Fos, AFT (CREB), and Maf families (33, 34). The transcriptional activity of Ap-1 is regulated by growth factors, cellular stress, and viral or bacterial infections. Both MAPKs and PI3K signaling pathways can regulate the abundance and transactivation of c-Jun and c-Fos (35) (Fig. 4A). To investigate this pathway, we used the PI3K and MAPK inhibitors LY294002 and U0126, respectively, on miR21 regulation. Northern blot analysis showed that inhibition of both the PI3K and MAPK pathways blocked miR21 upregulation in HeLa229 cells (Fig. 4B). In addition, chlamydial infection resulted in a strong phosphorylation of c-Jun and c-Fos, reflecting activation of the transcription factor Ap-1 (Fig. 4C,D). We also used a specific inhibitor of Ap-1, Sp100030, to rescue the effect (Fig. 4E). Ap-1 inhibition resulted in the depletion of miR21 level (Fig. 4F) which rescued the PTEN level and subsequently affected the bacterial growth (Fig. 4E). Our previous study has shown that *Chlamydia* infection results in consistent PI3K activation which leads to the depletion of TP53 (5). Interestingly the human genomic PTEN locus has a TP53 binding element directly upstream of the PTEN gene suggesting that PTEN is transcriptionally regulated by TP53 (36) (Fig. 4G). To investigate if TP53 can overcome the miR21-dependent downregulation of PTEN upon *Chlamydia* infection, we overexpressed TP53 in host cells (Fig. 4H). Expression of TP53 rescued the loss of PTEN and subsequently limited the growth of *Chlamydia* (Fig. 4H). This effect was also pronounced when we used etoposide, a topoisomerase inhibitor that stabilizes TP53 (Fig. 4I). Both these experiments revealed a feedback loop connecting miR21, PTEN, PI3K, and TP53 and showed its pivotal role in chlamydial infection.

**Figure 4.**
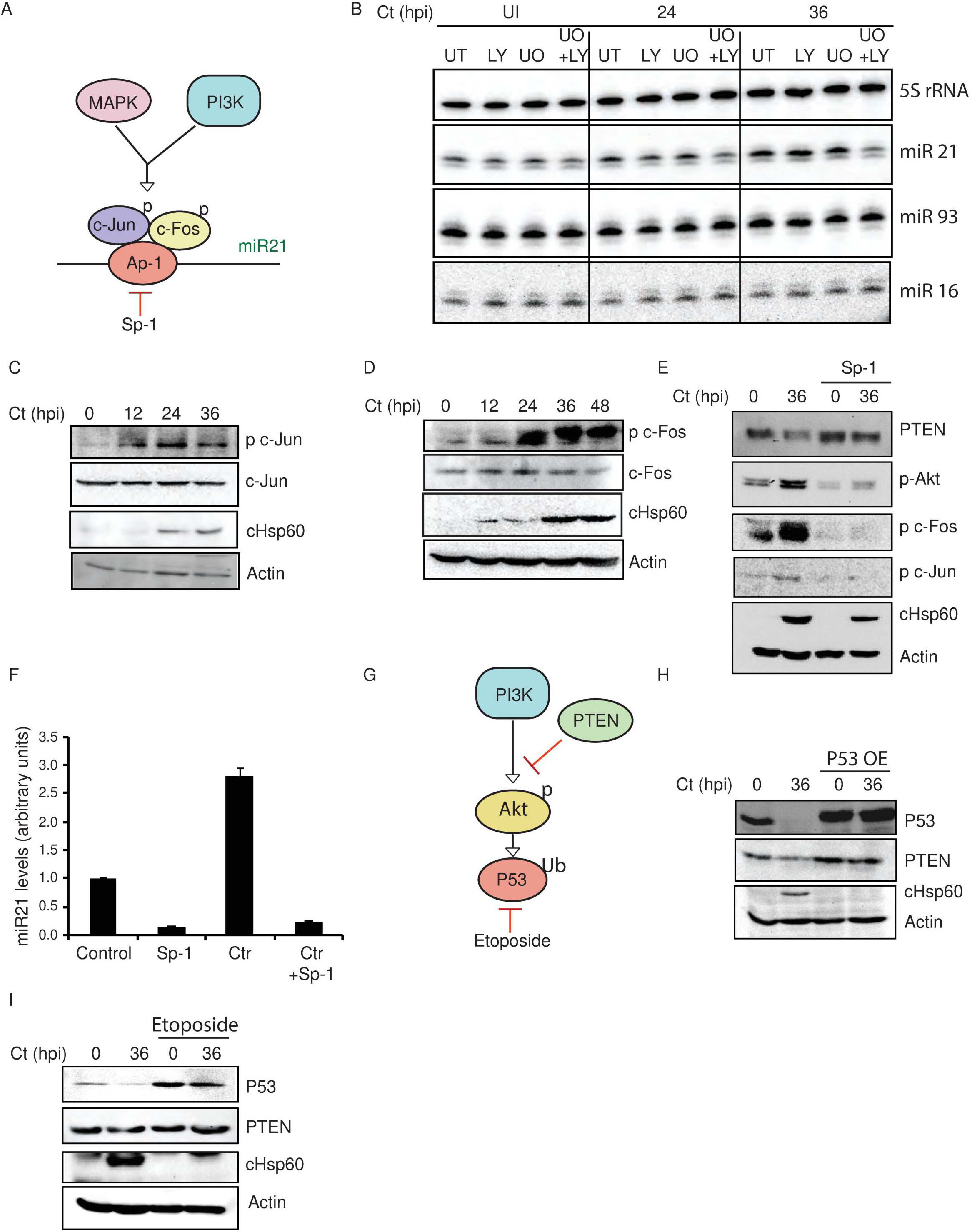
*Chlamydia trachomatis* regulates miR21 via MPAK and PI3K pathway**. A.** Cartoon showing molecular signaling pathway by which miR21 is regulated. **B**. HeLa229 cells were either left untreated or treated with PI3K inhibitor (LY294004) or MAPK inhibitor (UO126). Further these cells were either left uninfected or infected with Ct. The cells were analyzed using Northern blots to detect miR21 levels. miR93 and miR16 was used as a loading control. **C**. HeLa229 cells were infected with Ct for different time points, and the lysate was used for western blot analysis to detect p c-Jun and total c-Jun. n=3. **D**. the lysate from experiment **C** was used to detect the levels of c-Fos and p c-Fos. cHSP60 serve to detect Ct infection and beta actin serve as loading control. n=3. **E.** HeLa229 cells were either left untreated/uninfected or treated with SP-1 and infected with Ct. The cells were lysed for Western blot analysis to determine the levels of proteins. cHSP60 serve to detect Ct infection and beta actin serve as loading control. n=3. **F**. Control/Ct infected cells or SP-1 treated/Ct infected cells were taken for RNA extraction to determine the levels of miR21 via quantitative RT-PCR. n=3. **G.** Cartoon showing the role of PTEN in regulating the tumor suppressor p53. **H**/**I.** HeLa229 cells were either left untreated or over expressed with P53/infected or treated with etoposide/infected. The cells were lysed to analyze the levels of proteins. cHSP60 serve to detect Ct infection and beta actin serve as loading control. n=3.

### miR21 is required for the growth and development of *C. trachomatis in vivo*

All experiments in this study were initially performed in cell culture models. To determine whether PTEN, PI3K, and TP53 are similarly deregulated in a complex *in vivo* setting, we established a transcervical mouse infection model. Tissue lysates were prepared from infected reproductive tracts and analyzed for protein expression (Fig. 5A). Consistent with our *in vitro* findings, PTEN and TP53 levels were reduced in infected tissues, whereas total and phosphorylated Akt levels were increased.

**Figure 5.**
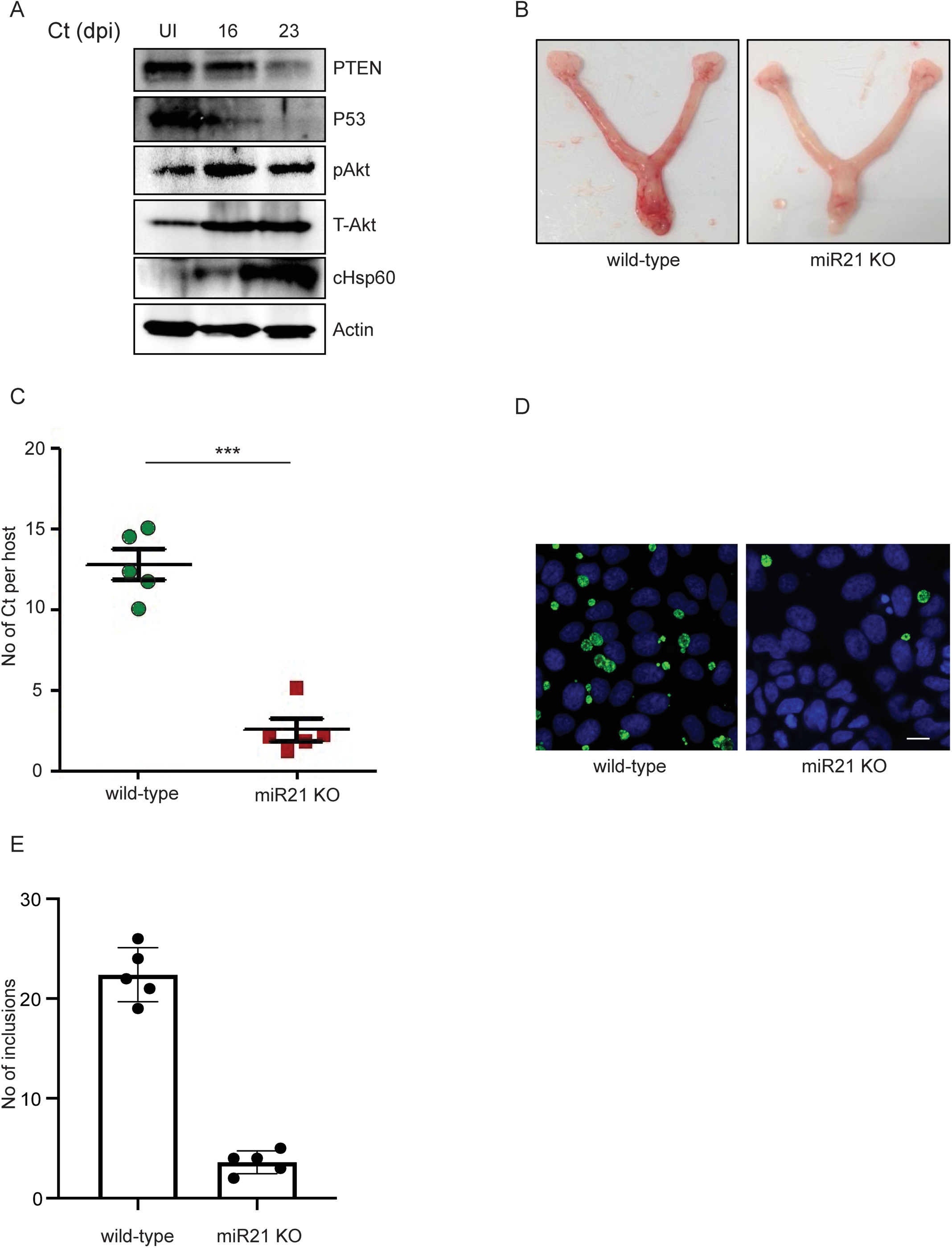
miR21 KO mice do not support chlamydial infection. **A**. The lysate from the control and infected uterine horns were subjected to western blot analysis to detect the levels of the respective proteins. cHSP60 serve to detect Ct infection and beta actin serve as loading control. n=3. **B.** The uterine horns of wild type and miR21 KO mouse infected with *C. trachomatis* for 21 days. n=5 **C**. Wild-type and miR21 KO mice were infected with *C. trachomatis* for 21 days. n=5. Total RNA was isolated and quantitative RT-PCR was performed to determine the copy number of the bacterium. Statistical analysis was performed using unpaired t test (t = 8.986, df=8). Error bar was defined as mean with ±SEM. n = 5, ***p ≤ 0.001. **D**. Progeny from infected mouse organoids: Organoids were derived from wild-type and miR21 KO mice and infected with *Chlamydia* for 6 days. Then, the organoids were lysed and released bacteria were used to infect freshly plated HeLa cells. The cells were fixed and stained for *Chlamydia* (Ct, green) and phalloidin (red) n = 3. Scale bar = 10 µm. Representative images of three independent experiments are shown. **E**. Number of inclusions from **D** was counted to plot the graph presented in scatter box plot overlaid with data point overlaid in the graph. p value was determined by two tailed unpaired *t* test (t = 14.33, df=8). Error bar was defined as mean with ±SEM. n = 5. **** indicates p value <0.0001.

To assess whether miR21 is required for *Chlamydia* infection and growth *in vivo*, C57BL/6 wild-type and miR21 knockout (KO) mice were infected for 21 days. Notably, infected miR21 KO mice did not develop detectable pathology, in contrast to infected wild-type mice (Fig. 5B). Quantification of bacterial burden in the female reproductive tract by RT-PCR revealed significantly reduced bacterial loads and impaired progeny formation in miR21 KO mice compared to wild-type controls (Fig. 5B).

The striking absence of pathology in infected miR21 KO mice suggested reduced inflammation, particularly diminished neutrophil activation, which is a major driver of pathology during *C. trachomatis* infection (37). In addition, several studies have linked PTEN to neutrophil function (38, 39). To test whether miR21 deficiency alters neutrophil bactericidal activity, we isolated neutrophils from the bone marrow of wild-type and miR21 KO mice and infected them *ex vivo* for 20 hours. Neutrophils were then lysed, and the recovered bacteria were used to infect HeLa cells to quantify infectious progeny, as previously described (40, 41). No differences in bacterial survival or progeny formation were observed between bacteria recovered from wild-type and miR21 KO neutrophils (Fig. S4), indicating that miR21 deficiency does not affect neutrophil-mediated killing.

To determine whether the reduced bacterial burden observed *in vivo* was due to defects in epithelial cells, we generated epithelial organoids from wild-type and miR21 KO mice and established infection (Fig. S3). Organoids derived from miR21 KO mice were noticeably smaller than those from wild-type mice (Fig. S3). Importantly, miR21 KO organoids failed to support bacterial growth, as demonstrated by immunostaining and infectivity assays performed on infected organoids (Fig. 5D,E).

Given that the miR21 target PTEN is a well-established tumor suppressor and regulator of glucose metabolism (42), we performed metabolomic analyses of uninfected and infected wild-type and miR21 KO organoids using mass spectrometry. While no major shifts in global metabolic pathways were observed, subtle regulatory differences between conditions were detected. These findings are consistent with previous proteomic analyses comparing wild-type and miR21 KO cell lines in cancer models. Together, these findings demonstrate that miR21 is a critical host factor that facilitates *C. trachomatis* pathogenesis *in vivo* by promoting epithelial permissiveness and associated signaling alterations.

## Discussion

*C. trachomatis* subverts multiple host signaling pathways to adapt to the intracellular niche.

Due to its reduced genome, the bacterium is heavily dependent on host-derived metabolites, including carbohydrates, amino acids, nucleotides, and lipids (9). This metabolic reliance necessitates extensive reprogramming of host cellular processes to sustain bacterial replication and survival.

The PI3K-pathway, one of the most frequently dysregulated signaling cascades in cancer, coordinates cellular nutrient uptake and utilization, regulating glucose, glutamine, lipids, and nucleotide metabolism to activate proliferation (43). *C. trachomatis* activates PI3K early during infection, and this activation persists throughout its developmental cycle, although the underlying mechanism remains incompletely defined. Sustained PI3K-Akt signaling likely ensures adequate metabolic support for bacteria growth while simultaneously protecting infected cells from apoptosis. Accumulating evidence further indicates that *Chlamydia* infection induces a tumor-like state in the host cell characterized by constitutive PI3K activation, depletion of TP53 (4, 5) and stabilization of proto-oncogene c-Myc (8). Elucidating the mechanisms responsible for persistent activation of this tumorigenic pathway in infected cells is therefore critical in understanding and controlling chlamydial pathogenesis, as well as the long-term pathological consequences associated with chronic infection.

Human cells tightly regulate PI3K activation to maintain signaling homeostasis. The tumor suppressor PTEN, a lipid phosphatase that specifically catalyzes the dephosphorylation of 3‘ phosphate of the inositol ring in PIP_3_, is a central negative regulator of the PI3K pathway. It is frequently inactivated in endometroid ovarian cancer. Our previous studies demonstrated that EphA2 functions as both an invasion and intracellular signaling receptor that recruits and activates the PI3K pathway during *C. trachomatis* infection (15). In addition, chlamydial effectors such as TepP have been shown to spatially regulate the activation of PI3K in the infected cells (16).

In the present study, we identified a mechanism by which the bacterium maintains the constitutive PI3K signaling through depletion of its endogenous inhibitor, PTEN. This work originated from a miRNA screen performed in infected host cells. Although previous studies have shown that infertility and caspase activation induced by *Chlamydia*-infection coincide with Dicer cleavage and altered microRNA expression in the oviducts of infertile mice, we did not observe depletion of Dicer levels in infected cells (Fig. S5). Interestingly, the miR21-dependent PTEN depletion emerged as a highly conserved response, observed across multiple different cell lines and mouse tissue infected with distinct *Chlamydia* serovars (Fig. 1, S1). Fimb cells may of particular interest since most ovarian cancers originate in the fimbriated end of the fallopian tube. The identification of miR21 also completes a regulatory feedback loop connecting AP-1 activation to PI3K signaling and unveils an additional molecular node connecting chlamydial infection with cancer-associated pathways. Importantly, independent of other regulatory inputs regulating PI3K and P53, inhibition of AP-1 abrogates PI3K activation and significantly impaired bacterial growth (Fig. 4) confirming previous results (44). These data position the AP-1-miR21-PTEN axis upstream of sustained PI3K signaling and highlights its potential as a therapeutic target for antibiotic-independent control of infection. This concept is further supported by the marked inhibition of chlamydial propagation in a miR21 KO mouse infection model (Fig. 5).

Previous work established a role of miR21 in macrophages phagocytosis during Listeria infection (45) and in macrophages and neutrophils during sepsis (46). We therefore investigated the neutrophil as the central myeloid cell in defence against *Chlamydia* infection as a possible target for the infection phenotype observed in the mouse model. However, there were no differences in survival and development of Ct in wildtype and KO neutrophils, indicating that miR21 has no major impact on the phagocytic and bactericidal activity in these cells likely because Ct does not effectively replicate within these immune cells and therefore has reduced metabolic requirements. Instead, we observed a growth defect of Ct in organoids derived from miR21 KO mice compared to organoids from WT mice, demonstrating the first phenotype of miR21 of bacterial infection in primary epithelial cells. Using WT or miR21 deficient mice and *in vitro* assays with mouse macrophages, Hackett et al. demonstrated that miR21 dampened glycolysis and ultimately decreased IL-1β production (47). Our attempts to detect differences in the metabolic response of infected versus non-infected organoids from WT and miR21 KO mice revealed multiple minor differences but no clear change in metabolic traits (data not shown). Further studies are required to determine the mechanism underlying of miR21-mediated Ct infection and inflammation.

In this study we focused on a single target of miR21; however, miR21 is known to regulate a broad network of genes. Its upregulation has been implicated in numerous cellular processes spanning virtually all hallmarks of oncogenesis, including proliferation, invasion, inflammation, metabolic reprogramming, angiogenesis, and evasion of cell death (48). In the context of infection, elevated levels of miR21 may therefore modulate multiple gene targets simultaneously, creating an interconnected signaling landscape that potentiates oncogenic pathways and promotes tumor initiation and progression.

## Methods

### Cell culture and *Chlamydia trachomatis* culture

HeLa229 (ATCC^®^CCL-2.1^™^), human Fimb: epithelial cells isolated from fimbriae of healthy donors after hysterectomy and Huvec cells (ATCC^®^ CRL-1730^™^) were used in the experiments. HeLa229 and human Fimb cells were grown in standard cell culture media (RPMI GlutaMax (Gibco) with 10% Fetal calf serum (FCS) (Sigma-Aldrich F7524)), heat-inactivated at 55°C for 30 min. Huvec cells were grown in Medium 200 (Gibco^TM^ M200500) containing 1x LSGS (Gibco^TM^ S00310).

*Chlamydia trachomatis* (serovar L_2_/434/Bu) (ATCC® VR-902B™) */ Chlamydia muridarium* (ATCC-VR-123) was cultured in HeLa229 cells as described before. Briefly HeLa229 cells were infected with Ct with a multiplicity of infection (MOI) of 1 for 48 hours (h) in 150 cm^2^ flasks. The cells were then scrapped with a rubber policeman and lysed with glass beads (Sigma, 5mm) for 3 minutes (min). The cell lysate was collected and centrifuged at 2000 revolutions per minute (RPM) for 10 min at 4°C to remove the cell debris. The supernatant was further centrifuged at 25000 RPM for 30 min at 4°C. The bacterial pellet was then washed with SPG buffer (0.25 M sucrose/10 mM sodium phosphate/5 nM glutamic acid) and passed through G30 and G18 syringes to break the clumps. The bacteria were further aliquoted and frozen at-80°C until used. Mycoplasma contamination in cell culture and bacterial preparation was routinely checked and controlled via PCR.

### Cloning, constructs used, and transfection in cells

The plasmids used in the study are described in the supplementary file. Briefly, miR21-mimic RNA, scrambled RNA, pCDNA3 expressing miR21 (miR21OE), pCDNA3 expressing miR21 fused with GFP-PEST (miR21 GFP PEST), or pCDNA3 expressing GFP-PEST (GFP-PEST), pCDNA3 expressing PTEN 3’UTR and pCDNA3 expressing PTEN 3’UTR with point mutation were cloned and used in the study. Cells were transfected with plasmid DNA at a confluency of 60% with X-tremeGENE^TM^ HP DNA transfection reagent (Roche) and OptiMEM transfection medium (Gibco) in 5% FCS medium. After 5 h, the transfection medium was replaced by fresh RPMI supplemented with 5% FCS. The cells were infected with Ct after 24 h of transfection and after 24-36 hpi, the cells were taken for analysis.

### Inhibiting miRNA activity

For inhibition of miRNA activity in cultured cells, antagomirs (chemically modified antisense oligonucleotides complementary to the mature miRNA) were delivered using a lipid-based transfection method. Briefly, cells were seeded to reach approximately 60–70% confluency at the time of transfection. The antagomir (commonly used at a final concentration of 25–100 nM, with 50 nM as a starting point) was diluted in serum-free medium such as Opti-MEM, and the transfection reagent was diluted separately in the same medium. After a short incubation (5 minutes), the two solutions were combined and allowed to form complexes for 15–20 minutes at room temperature. The complexes were then added dropwise to cells maintained in antibiotic-free complete medium. Cells were incubated for 24–72 hours before downstream analysis. Successful inhibition was validated by assessing the levels of miRNA and its targets at the mRNA (qRT-PCR) and protein (Western blot) levels, alongside appropriate scrambled and mock-transfected controls.

### Western blotting and Immunostaining

Western blotting was performed as explained elsewhere(10). Lysates for Western blot analysis were prepared by directly lysing cells in SDS sample buffer (62.5mM Tris, pH 6.8, 2% SDS, 20% glycerol, and 5% ß-mercaptoethanol) in ice. Briefly, protein samples were separated in the 6-12% SDS-PAGE (Peqlab) and transferred to a PVDF membrane (Roche) in a semidry electroblotter (Thermo Fischer Scientific). The membrane was further blocked in tris buffer saline containing 0.05% Tween20 and 5% bovine serum albumin or dry milk powder. The primary antibody against PTEN (ab137337) was purchased from Abcam. The T-ERK (cs-9180), pERK (cs-9106), T-AKT (cs-9272), pAKT Ser473 (cs-9271), T-MEK (cs-9122) and pMEK (cs-9121), Dicer, c-Jun, p c-Jun, Jnk, pJnk, c-Fos, p c-Fos, p53, and GFP were obtained from Cell Signaling. Chlamydial HSP60 (sc-57840) was purchased from Santa Cruz Bioscience and ß Actin antibody from Sigma (A5441). Proteins were detected with secondary antibodies coupled with HPR (Santa Cruz Bioscience) using the ECL system (Pierce) and Intas Chem HR 16-3200 reader. Quantification of blots was done by FIJI (ImageJ) software.

### RNA isolation cDNA isolation and qRT-PCR

Cells were either left uninfected or infected with Ct at a MOI 1 for respective time points. The RNA was isolated using a RNeasy mini kit (Qiagen) as indicated by the manufacturer. The RNA was eluted in ultrapure water and was stored at-80°C until used. 1μg of RNA was used for cDNA synthesis and was performed using the RevertAid RT kit (Thermo Fisher Scientific) and Perfecta SYBR Green FastMix (Quanta Biosciences). Primers for qRT-PCR were as follows: GAPDH-forward, 5′-CGTCTTCACCACCATGGAGAAGGC-3′, and reverse, 5′-AAGGCCATGCCAGTGAGCTTCCC-3′, PTEN-forward, 5′-CCAGGACCAGAGGAAACCT-3′, and reverse, 5′-GCTAGCCTCTGGATTTGA-3′.

For miRNA, the cDNA preparation and qRT-PCR for miRNA were performed using the miScript II RT kit and miScript SYBR Green PCR kit (QIAGEN) according to the manufacturer’s protocols. Custom-made primers for all the miRNAs studied were also bought from QIAGEN. qRT-PCR of miRNAs was performed using the miScript PCR System (QIAGEN) or miRCURY LNA universal RT miRNA PCR (Exiqon) with specific forward primers or LNA primer sets for individual miRNAs (QIAGEN or Exiqon) and endogenous control, U6 SnRNA (QIAGEN or Exiqon). All experiments were performed on a StepOnePlus real-time PCR platform (Applied Biosystems) according to the manufacturer’s protocol. Data were analyzed using StepOne Software v2.3 and GraphPad Prism7.

### Northern blotting

Northern blots for miR21 were performed using oligonucleotide probes for miR21-5p (5-TCAACATCAGTCTGATAAGCTA-3′) and U6 snRNA (5-CACGAATTTGCGTGTCATCCTT-3′) according to the protocol described by the Narry Kim Lab (http://www.narrykim.org/en/protocols). U6 snRNA was used as a loading control for all experiments done in HUVECs. The chlamydial small RNA described as CtrR5 (49) was used as a marker for *C. trachomatis* infection (5′-CAGCACCCCTCTGAGTTCTCCC-3′). Hybond-XL (GE Healthcare) nylon membranes were used for Northern blots. Decade Markers System from Thermo Fisher Scientific was used to radiolabel RNA ladder for Northern blots.

### Fluorescent assisted cell sorting (FACS)

Hela cells were seeded in six-well plates for 12 h and transfected with Empty-GFP-PEST or GFP-PEST-miR21 for 12 h before infection with Ct. At the endpoint, cells were trypsinized and suspended in a cell-specific medium. The fluorescence of GFP and dsRED were measured simultaneously in FITC (509 nm) and PE (560 nm) channels by flow cytometry using a BD FACS Aria III. The cells were gated for increasing fluorescence in the FITC channel and normalized to the PE channel for variations in plasmid expression.

### Infectivity assay

Cells were left untreated or treated with inhibitors/ transfected with the respective constructs. The cells were infected with *C. trachomatis* for 36 hpi and washed once with PBS. The cells were then lysed manually by glass beads. Different dilutions of the lysed sample with infectious EBs were used to infect freshly plated HeLa cells, 24 hpi the cells were analyzed using immunostaining (counting the inclusion forming units) or immunoblotting.

### Transcervical mouse infections and determination of bacterial burden

C57BL6 and miR21 KO mice were used for the experiments. All animal experiments were performed by protocols approved by the animal care and experimentation of German Animal Protection Law approved under the Animal (Scientific Procedures) Act 1986 (project license 55.2-2532-2-762). The mice used for the experiment were between 10-14 weeks old. Five days before transcervical infection, mice were treated subcutaneously with 2.5 mg of DepoProvera (medroxyprogesterone acetate). The mouse infection and determining the bacterial burden were performed as published before. Data were analyzed using the Step One Plus software package (Applied Biosystems) and expressed as the ratio of chlamydial genome to host genome (*lytA/*synectin). GraphPad Prism 7 was used to generate a scatter column chart and perform statistical analysis. One-way analysis of variance (ANOVA) was performed with the significance level set to less than 0.01. Statistical analysis was performed to decide the sample size used in mouse infection by the Institute of Mathematics, University of Würzburg under the allowance A2 55.5-2531.01-49/12. All mouse experiments were carried out with five female mice per treatment group. Mice in each experiment were age-matched and cage mates were randomly distributed into different treatment groups to avoid cage effects.

### Organoid generation and bacterial infection

Mouse fallopian tube samples were processed within 2 h the mouse was sacrificed. Briefly, the tissue samples were washed with DPBS (Gibco) and placed in a sterile Petri dish where the samples were minced into small pieces using a sterile scalpel. The tissue samples were incubated with collagenase and dispase for 1 h at 37°C before being placed between two glass slides and pressed gently. Further, the tissue samples were suspended in RPMI and centrifuged at 1000 RPM for 10 minutes. The cell pellet was resuspended with Matrigel (Corning) and placed as 50 µl drops in the center of a 24-well plate. The Matrigel was solidified for 20 min at 37C and further incubated with 500µl of prewarmed enriched media (DMEM advanced (Sigma), Wnt media (obtained from the supernatant of cells (ATCC) 25%), R-Spondin (25%), Noggin (10%), B27 (Thermo Scientific), Penicillin/streptomycin (pen/Step) (1%), Nicotinamide (Sigma, 1 mM), Mouse EGF (50ng/ml) (Thermo Scientific), Human FGF (100ng/ml) (Thermo Scientific), TGF inhibitor (0.5µM) (Tocris), ROCK inhibitor (10µM) (AbMole Biosciences)). After the organoids were established pen/ strep was removed from the media and infection was performed. The organoids were visualized for infection rate using immunostaining.

### Metabolic profiling

C57BL6 and miR21 KO organoids were seeded in a 6 well-plate and left uninfected or infected with *C. trachomatis* MOI 1. The medium was collected after incubation for 30 h and snap-frozen in liquid nitrogen and the cells were washed with ice-cold 154 mM ammonium acetate (Sigma) and snap-frozen in liquid nitrogen. The cells were harvested after adding 480 μl cold MeOH in H2 O (80/20, vol/vol; Merck) to each sample containing lamivudine standard (10 μM; Sigma). The cell suspension was collected by centrifugation and transferred to an activated (by elution with 1 ml CH 3CN (Merck)) and equilibrated (by elution with 1 ml MeOH in H2O (80/20, vol/vol)) RP18 SPE-column (Phenomenex). The eluate was collected and evaporated in a SpeedVac concentrator. The residue was dissolved in 50 μl of 5 mM NH4OAc (in CH3CN and H2O; 25/75, vol/vol). Each sample was diluted 1:2 (cells) or 1:5 (medium) in CH 3 CN. The sample (5 μl) was applied to a HILIC column (Acclaim mixed-mode HILIC-1, 3 μm, 2.1 × 150 mm). The metabolites were separated at 30 °C by liquid chromatography using a DIONEX Ultimate 3000 UPLC system (Solvent A: 5 mM NH4 OAc in CH3 CN and H2 O (5/95, vol/vol); Solvent B: 5 mM NH4OAc in CH3CN and H2O (95/5, vol/vol); gradient: linear from 100% Solvent B to 50% Solvent B in 6 min, followed by 15 min of constant 40% Solvent B). Mass spectrometry analysis was conducted on a Thermo Scientific QExactive instrument in alternating positive and negative modes. Peak determination and semi-quantitation were performed using the TraceFinder software. Data was normalization of total metabolites. MetaboAnalyst 5.0 was used to analyze data.

### Neutrophil isolation of mice bone marrow and infection

Mouse bones of the lower extremities were cleaned of remaining muscle tissue and rinsed with 1x PBS in a petri dish on ice. With a scalpel the epiphysis was cut off from the bones, and a syringe with a 25G needle was used to flush the bone marrow cells with 1x PBS through a 100 µm cell strainer into a conical tube. Cells were collected via centrifugation at 427 x g for 7 min at 4°C. Pellet was resuspended in 0.2% NaCl and followed by addition of 1.6% NaCl equal volume to lyse red blood cells. To collect bone marrow cells, cells were centrifuged at 427 x g for 7 min at 4°C. Pellet was resuspended in ice cold 1x PBS and purified via a Histopaque density gradient. Histopaque 1119 was overlayed with Histopaque 1077 and cell suspension on top, centrifuged at 872 x g for 30 min at room temperature and without break. Neutrophils were collected at the interface of the Histopaque layers. Purity was analysed via flow cytometry and viability by trypan blue addition while determine neutrophil counts. Equal amounts of C57BL6 and miR21 KO neutrophils (5 x 10^5^ cells in 500 µl RPMI w/o serum) were taken and infected with *C. trachomatis* MOI 10. After 20 h of infection, neutrophils were lysed with water, centrifuged at 188 x g for 5 min at room temperature, and supernatant was added to freshly seeded HeLa cells. 24 hpi inclusions were counted, and inclusion per cell ratio determined. Analysis was done with GraphPad Prism.

## Author contribution

KR and TR designed the experiments, KR, SRC, MA, and NV performed experiments. JW provided cell lines. KR and TR wrote the manuscript.

## Supporting information

Plasmids and primers

## Acknowledgment

We thank Dr. Ana Eulalio and Malavika Saran for important support with miR-seq. We thank Eric Olson (UT Southwestern Medical Center at Dallas, USA) for the miR21 KO mice. This work has been funded by the European Research Council grant to T.R. (ERC-2018-ADG/NCI-CAD), KR (Frauenbüro) and SRC (Graduate School of Life Sciences) received additional funding from the University of Würzburg.

---

**Extended figure 1.**
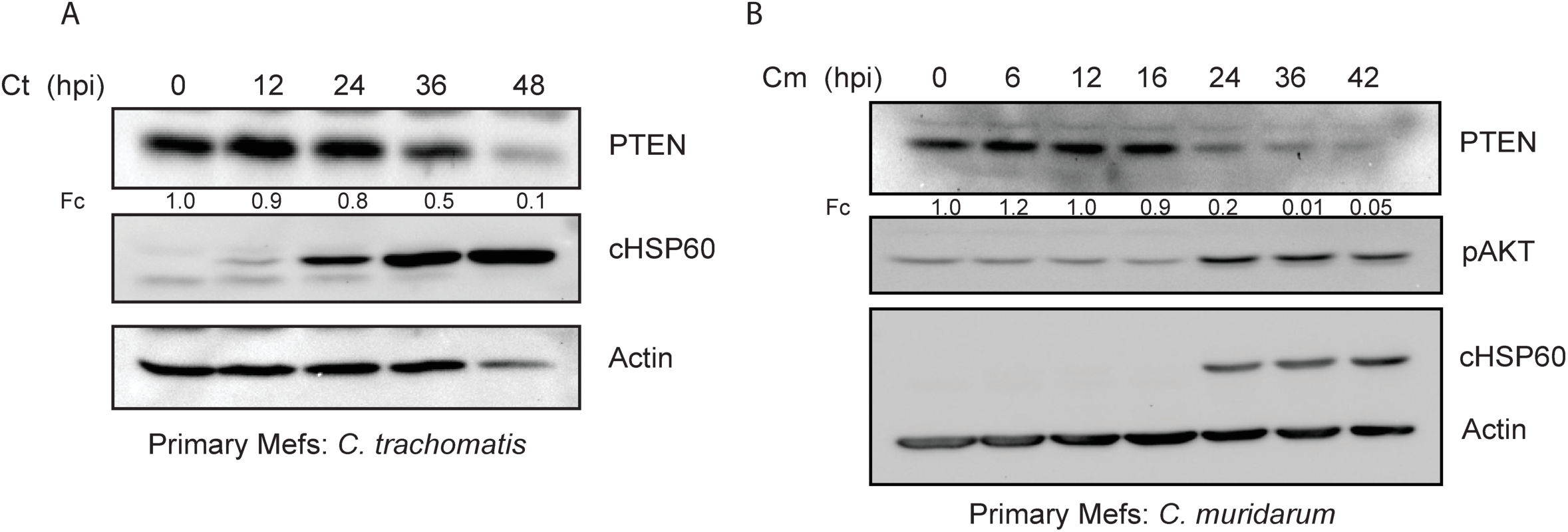
**A.** Primary mouse embryonic fibroblasts (Mefs) were infected with Ct at a MOI 1 for different time points. The cells were lysed and analyzed via western blotting. (n=3). **B.** Primary Mefs were infected with *Chlamydia muridarum* (Cm) for different time points. The cells were lysed and analyzed via western blotting. (n=3).

**Extended figure 2.**
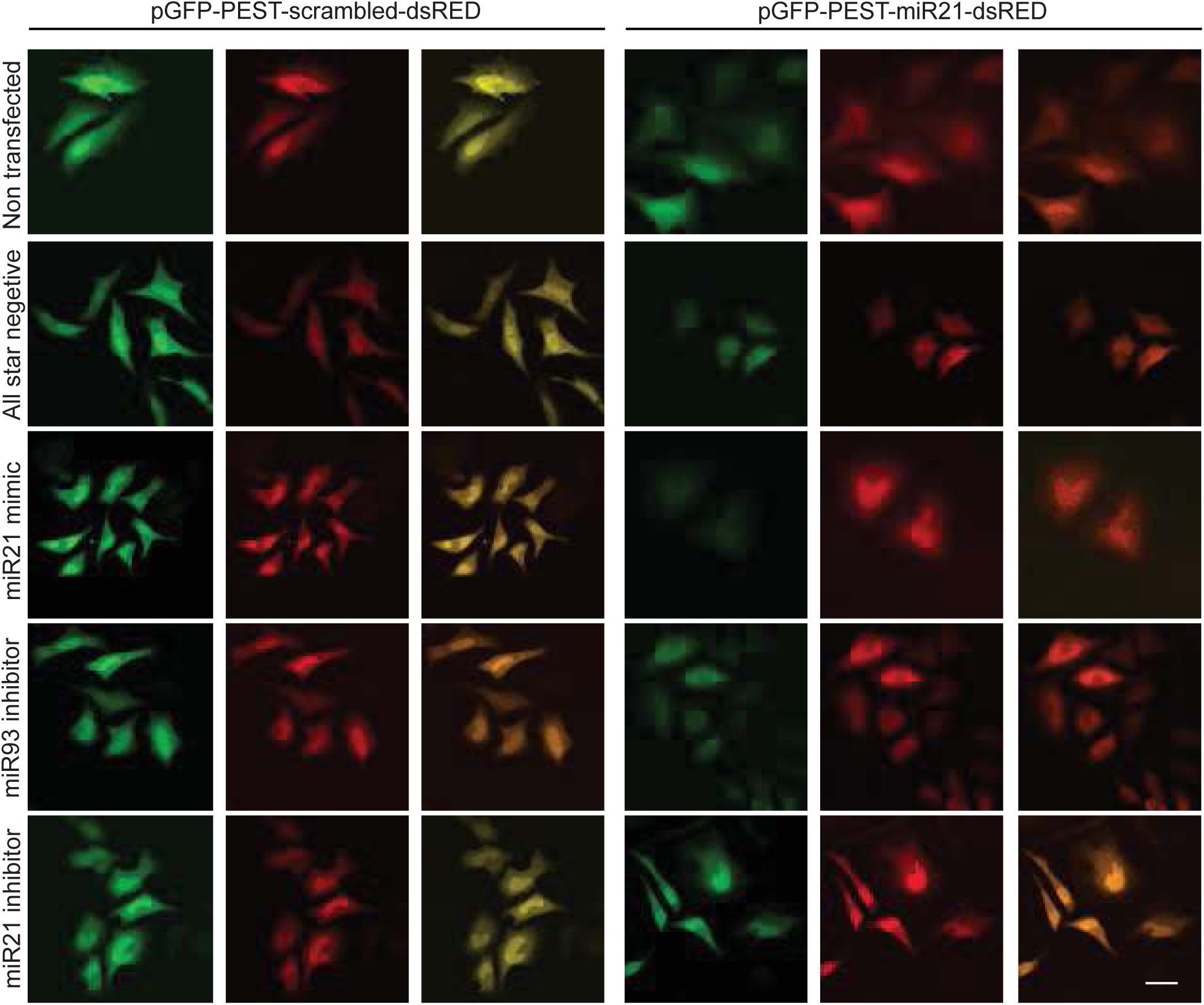
PEST-GFP scrambled dsRED construct with scrambled or miR21 binding site (PEST-GFP miR21 dsRED) were transfected in HeLa229 cells. The cells were further, either left non transfected or transfected with All start negative (scrambled peptide), miR21 mimic, miR93 inhibitor and miR21 inhibitor. The cells were fixed and analyzed using immunostaining. Green indicates GFP signal, red-dsRED. The images represent three independent experiments. (n=3).

**Extended figure 3.**
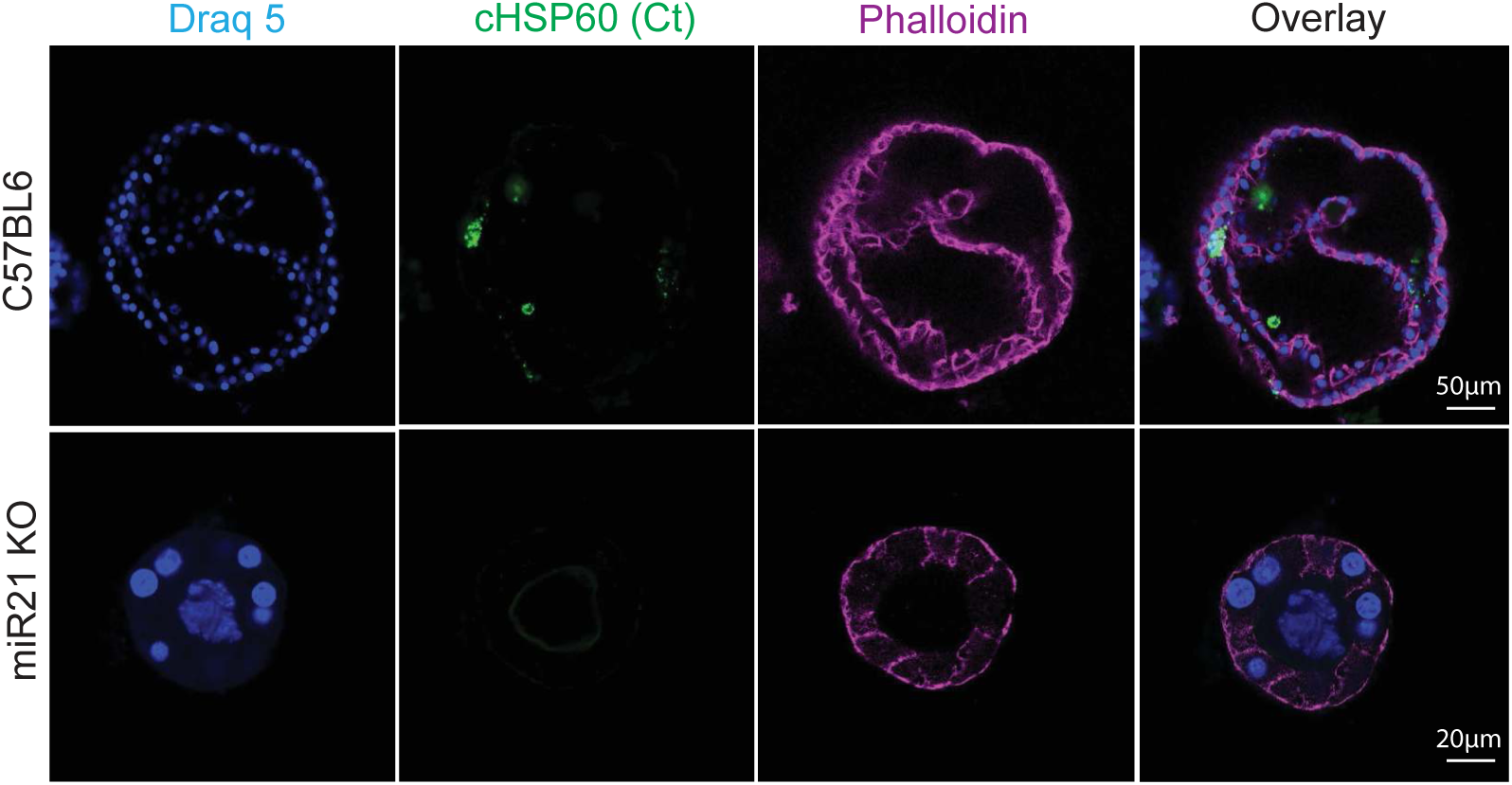
Organoids were derived from C57BL6 and miR21 KO mice and infected with *Chlamydia* for 6 days. The organoids were fixed and stained for *Chlamydia* (Ct, green), nucleus (blue, Draq5) and phalloidin (red) n = 3. Representative images of three independent experiments are shown.

**Extended figure 4.**
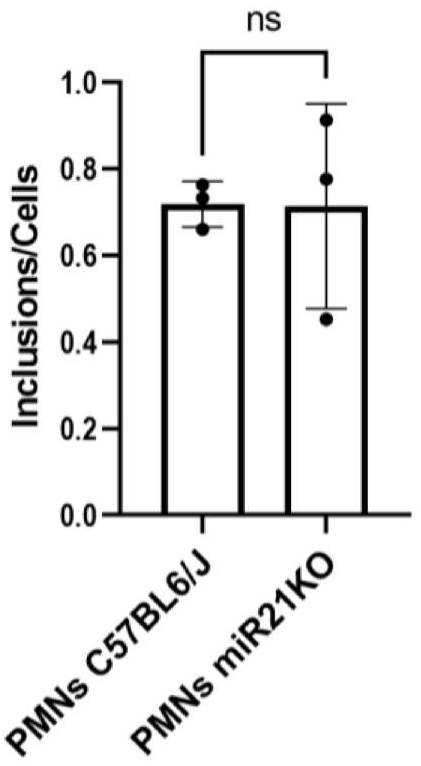
Bone marrow–derived neutrophils isolated from C57BL/6 wild-type and miR21 knockout (KO) mice were infected with *Chlamydia* for 20 h. Following infection, neutrophils were lysed and the recovered bacteria were used to infect HeLa cells for 24 h. Chlamydial inclusions were quantified and are presented as the inclusion-per-cell ratio (n = 3).

**Extended figure 5.**
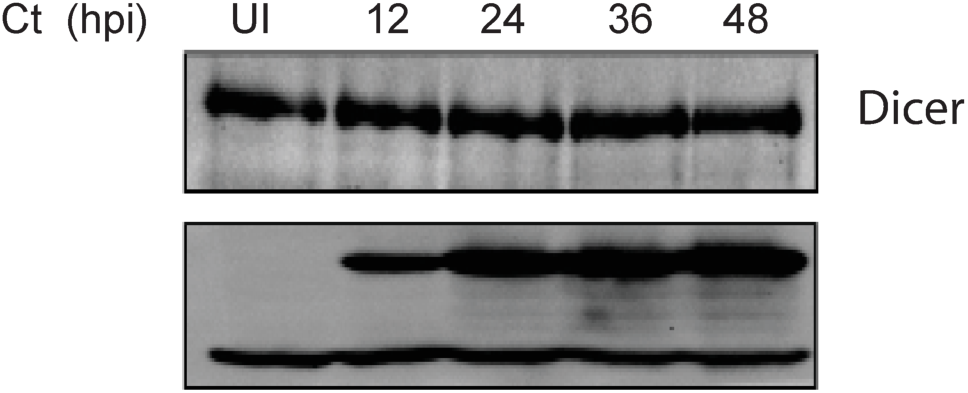
HeLa229 were infected with Ct at a MOI 1 at different time points. The cells were lysed and analyzed via western blotting. (n=3).

