## Supplementary material for "*Chlamydia trachomatis* infection upregulates MicroRNA21 to deplete tumor suppressor PTEN": Plasmids and primers

**Supplementary File**

| **Sl No** | **Plasmids** | **Primers used** |
| --- | --- | --- |
| 1. | pCDNA3- miR21 | Fwd: TATTTATTTAGTTATGACTGTGACGAC  Rev: TTTGCCTGGTAGGAAAATAAACAG |
| 2. | pCDNA3- Scrambled | Fwd: GATCGTACGGCTTAGGCTATACGG  Rev: GGTACCGTAGGCTGAGTCGTCA |
| 3. | pcDNA3- miR21 GFP PEST | Fwd: CGGCGGTAGCTTATCAGACTGATG  Rev: CCAGTCGAGGGTCCGAGGTATT |
| 4. | pcDNA3- GFP-PEST | Fwd: CCGACAACCACTACCTGAG  Rev: CCTCCGAATGAGAGTGTTTCG |
| 5. | pcDNA3- PTEN 3'UTR (PUG) | Fwd: ATCGGCGGCCGCC  Rev: ACACCTCTAGACGAA |
| 6. | pcDNA3- PTEN 3'UTR SDM (M-PUG) | Fwd: TGTGGCAACAGGTAAGTTTGCAG  Rev: CTGCAAACTTACCTGTTGCCACA |
